# Using the Concepts of Time-delayed Feedback Control in Biofeedback Systems in Children with ADD: A Preliminary Study

**DOI:** 10.1101/709469

**Authors:** Golnaz Baghdadi, Ateyeh Soroush, Farzad Towhidkhah, Reza Rostami

## Abstract

The reported efficacy of biofeedback is not consistent across studies. One issue that has not been addressed in previous studies is the effect of the reference signal in a biofeedback system.

We compared the effects of two different reference signals using a computational model and human experiments. One reference signal was fixed, and the other was variable and was designed based on time-delayed feedback control concepts. The computational model consists of a plant and a controller. Groups of typically developing and children with attention deficit disorder (ADD) participated in our experiments.

The computational model showed that time-delayed reference feedback control could have more positive effects on the performance than using a fixed reference signal. The model prediction was consistent with the results obtained from the experiments performed on children with ADD. However, in typically developing children, using time-delayed reference feedback control had a negative effect. Our preliminary results, which need to be investigated on more subjects, indicate the importance of selecting the reference signal in biofeedback systems according to the characteristics of each participant. The results remind the importance of “single-case methodology” in neurocognitive rehabilitation systems. Using computational concepts, some other predictions have also been provided, which need more investigations.

## 1. Introduction

Biofeedback is a rehabilitation tool [1]. In biofeedback therapy, a signal such as behavioral indices (e.g. response time or accuracy) [2], electroencephalography (EEG) [3], electromyography (EMG) [4, 5], trunk kinematic [6], speech-volume level [7], heart rate variability (HRV) [8], respiration rate, or the skin temperature [9] is recorded from individuals. Then, one or several features have been calculated from the signal. The features’ value is compared with a reference signal [10]. The result of the comparison is given back to the individual for modifying the performance [11].

Several brain areas are involved in processing internal or external feedbacks and modifying the neural and behavioral control policy based on the received feedbacks [12-14]. Internal and external feedbacks can cause a pleasant or unpleasant feeling in the person. These feelings play an important role in learning and functional policy of individuals [15]. Amygdala is involved in emotion processing and pleasure perception. Hippocampus keeps the memory of a pleasant experience. Prefrontal cortex and insula weight and allocate an appropriate amount of attention to the feeling receives from feedbacks. Some regions in basal ganglia are responsible for releasing Dopamine, which is an important neurotransmitter to encode the reward or punishment feedbacks [16, 17]. Beside basal ganglia, cerebellum processes the incoming sensory information and modulates them for sending to the motor cortex [18]. The anterior cingulate cortex is another brain region that detects the expectancy of feedback [12]. Some loops between the cortex and the thalamus are important in neurofeedback (NF) learning [19].

There is a coupling between the reward and attention circuits [20]. The impairment of reward processing has been reported in children with attention deficit hyperactivity disorder (ADHD) [21, 22], which is one of the neurodevelopmental disorders. The ADHD symptoms start from childhood, and some of them usually last into adulthood [23]. According to the DSM-V criteria, ADHD is categorized into three subtypes: 1-inattentive subtype, which is usually called attention deficit disorder (ADD); 2-hyperactive or impulsive subtype; and 3-combined subtype [24]. Neuroimaging studies associate the ADHD symptoms with small and low activity of the frontal lobes and other encephalon regions [23]. Common treatments for ADHD symptoms are medical treatments, behavioral therapy, or using some tools such as biofeedback.

Although the positive effects of feedback on cognitive functions and ADHD symptoms have been reported in many studies [1, 23, 25-28], there are number of research that show the helplessness or negative impact of biofeedback [29-31]. Ineffectiveness of feedback has been attributed to the working memory capacity, task demand, individuals’ ability, or the feedback type (i.e., output feedback, strategy feedback, positive or negative feedback) [29-32].

The main concern that has been followed in different studies is the selection of an appropriate feature or feedback type. In our knowledge, there is no study about the effect of the reference signal. The reference usually is an ideal value that is usually measured from the subject in an ideal condition or a normative population. Our main goal was to investigate the effect of the reference signal on the performance of children in an attention-demanding task.

Children with ADD usually become tired and lose their concentration sooner than typically developing children in tasks required sustained attention. Due to this problem, they have a higher average and variability of reaction time (RT) in comparison with typically developing children during a continuous performance test (CPT). In other words, their performance drops considerably because of the mentioned early fatigue. This early loss of concentration may happen because of an early change in their neural system. If a process can postpone this fatigue point, it can be applied as a cognitive rehabilitation method. In mathematical sciences, the point at which the behavior of a system changes, is called “bifurcation” point [33]. In that, the characteristics of the system’s behavior change by the alteration of a parameter in the system. This parameter is called “control parameter.” Accordingly, it can be concluded that the characteristics of the neural system of people with ADD bifurcate sooner than the typically developing ones. Time can be considered as the control parameter. Therefore, the goal is changing the time that people with ADD show signs of fatigue. For instance, if the concentration of people with ADD drops after about K minutes, our goal is replacing it with K+K’ minutes.

Previous studies showed the positive effect of using a feedback control algorithm in biological systems [34, 35]. Time-delayed feedback control is a method that is used to replace the bifurcation point in the science of control [36, 37]. In this control method, the controller of the system receives the difference between the current and the T steps ago output values of the system. Based on this difference, the controller applied a policy to equalize the mentioned output values. It has been shown that using such a mechanism can control the bifurcation point of the system. Therefore, designing a reference signal based on this controlling method may affect the impact of biofeedback therapy.

We investigated the effect of two different reference signals using a computational model and on the human performance experimentally. We first designed a black box computational model that could produce a pattern of RT variations. Then, we used a controller that could alter the pattern of RTs. The behavior of the designed system was assessed based on three conditions: without feedback, feedback with a fixed reference, and feedback with a variable reference that has been selected based on the time-delayed feedback control concepts. Then, three human experiments were designed to evaluate the computational model’s results. Two groups of typically developing and children with ADD did the experiments. In two conditions of experiments, children received visual feedback according to the pattern of their responses. Previous studies showed the positive effect of visual feedback on attentional performance [38]. The outcomes of this study indicated that the selection of the reference signal could affect the output of biofeedback therapy. However, more investigations are needed to determine an appropriate reference signal.

## 2. Method

### 2.1 Modeling Procedure

As mentioned, RT is an index that the pattern of its variability during a CPT is used as a criterion to diagnose ADD. A series of RTs recorded from an individual during a CPT looks like a random signal that its average and variability increases over time due to the lapse of attention. Fig. 1 shows RTs that have been recorded from two typically developing and ADD groups during a CPT.

**Fig. 1.**
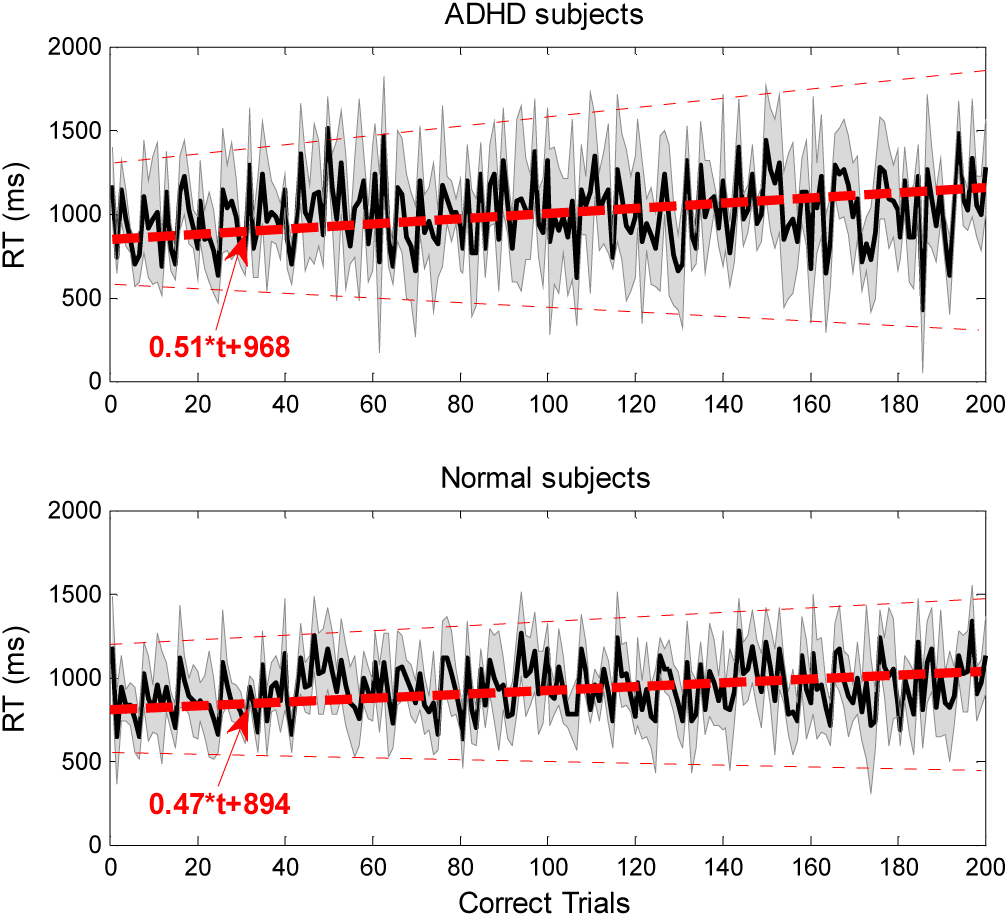
The changes of correct RTs (i.e., successful responses to the target) during a CPT in ADD (top plot) and typically developing (bottom plot) children. Thick dashed lines show polynomial fitted curves to the average of RTs in each group. Thicker solid lines indicate the average of RTs across the participants, and the shaded area shows between-subjects variability.

According to Fig. 1 and previous studies [39], it can be concluded that the mean of RTs increases from the first to the end of the CPT, the slope of this increment in children with ADD is higher than that of in typically developing children, because children with ADD get tired sooner during a task.

Equation (1) shows our suggested mathematical model to produce a signal that its pattern looks like the alteration of RTs over time.

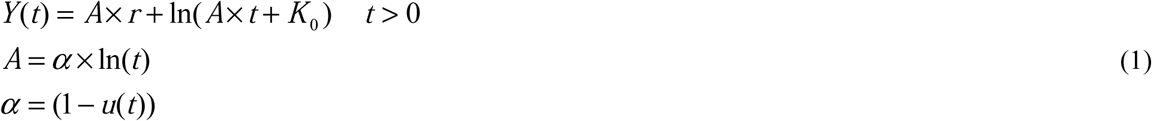

In this equation, *Y* is the output (i.e., RT signal), *u* is the input signal that can change from zero to one and affect the output pattern, *t* is the time sample (*t* > 0), *r* is a random number between zero and one, *K*_*0*_ affects the initial value of the output, *A* is a coefficient that regulates the growth rate of RTs. The growth rate depends on the value of *α*. The *α* value depends on the input value, *u*(*t*). Fig. 2 shows the output of the suggested model for two values of *u*. In Fig. 2, the value of *K*_*0*_ has been set on 20. This value has been selected arbitrarily; however, its alteration cannot change the pattern of the results. Changing the value of *K*_*0*_ only moves the graph up or down on the vertical axis.

**Fig. 2.**
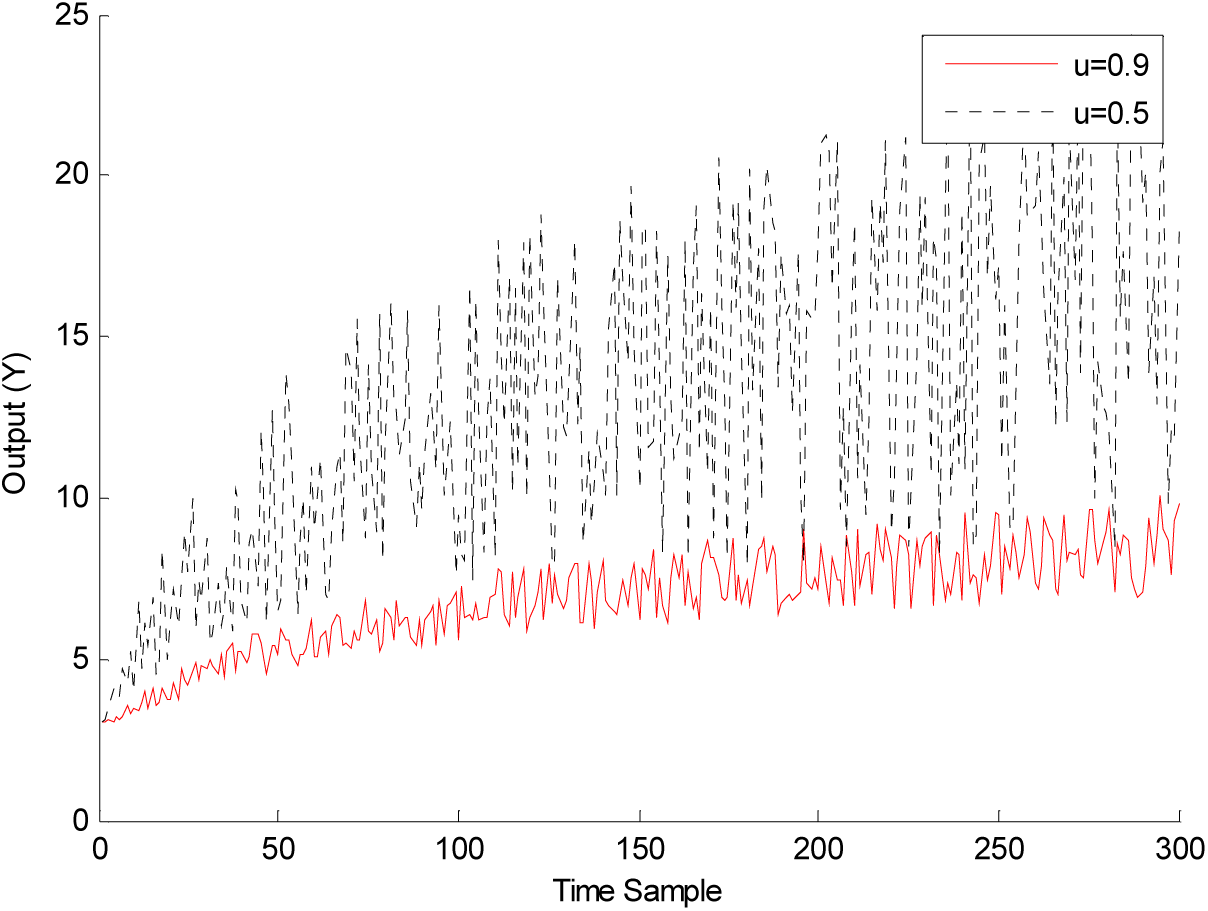
The output of Eq. (1) for two values of *u* and *K*_*0*_=20 (there is no feedback).

We considered some parts of the brain that the delay of their responses leads to different patterns of RT as a black box model. This model consists of a plant, (1), and a controller. As mentioned, the output of the plant, *Y*(*t*), is the RT signal. The input, *u*(*t*), is the controlling signal that can change the pattern of RTs (Fig. 2).

Decreasing the slope of the changes of RTs or the ratio of last to the first RTs can be considered as a control object to modify the output performance. The controller is responsible to produce an optimum controlling signal that leads to an appropriate RT. Its optimization procedure is based on the error between the output (*Y*(*t*)) and the desired RT, which is called the reference signal. Fig. 3 shows the components of the model.

**Fig. 3.**
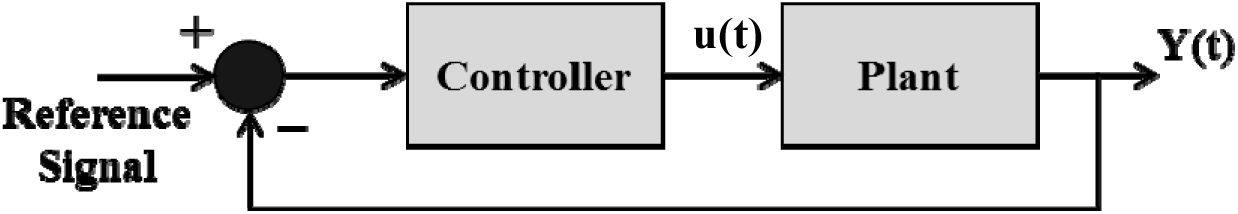
The components of the black box model with a desired fixed reference signal.

In the optimization procedure, the controller finds an optimum control signal (*u*) that causes the plant produces an RT that is near to the reference signal. Equation (2) shows the cost function that is minimized in the optimization procedure.

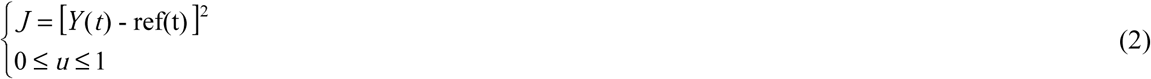

To find the optimum value of *u* that can minimize the cost function, Eq. (3) needs to be solved.

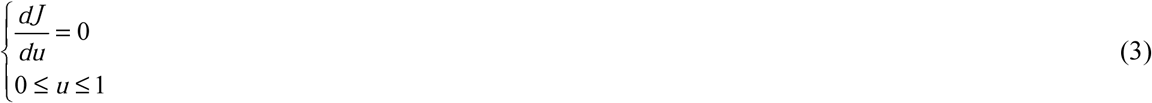

Substituting Eq. (1) in Eq. (2) and taking the mentioned derivative, Eq. (4) is obtained.

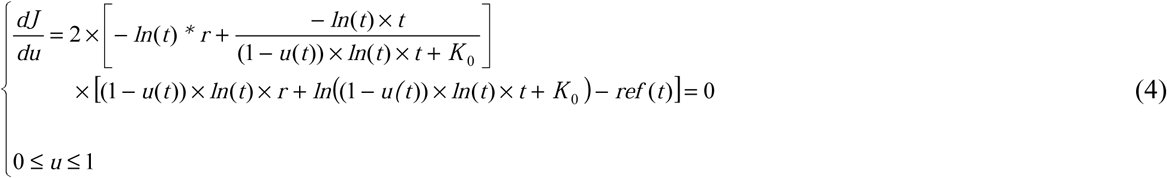

Equation (4) is solved using recursive methods.

In human experiments, the reference signal is defined internally [40] or externally [41]. In our model, we have focused on external feedback, which is called biofeedback in neurocognitive studies.

We investigated the effect of different reference signals in a biofeedback system. The signal that was given back to the model is the RT value in each sample of time. The performance of the plant output was investigated in three conditions:

1. Without feedback
2. With a desired fixed reference signal
3. With a variable reference signal designed based on the time-delayed feedback control concepts

In condition 2, the fixed reference signal was a mean RT that was calculated from a number of first correct trials (i.e., successful responses to the target). In this case, the controller tried to keep the output of the plant near the fixed reference over time by sending an appropriate control signal. However, due to some limitations, this goal might not be fully met. For example, the value of the control signal had limitations that did not allow the full compensation for the error between the output and the desired reference signal.

In time-delayed feedback control, the reference signal was the output of the plant at T step ago. Therefore, the goal of the controller was to keep the output of the plant near its value at T step ago. The value of T usually selected by the period of the plant. As shown in Figs. 1 and 2, the output of the plant in our model looks like a random signal. When the output of the plant is random, T is better to be set on one. Therefore, in condition 3, the reference signal was the output of the system with one sample delay. Fig. 4 shows the model design based on time-delayed feedback control concepts.

**Fig. 4.**
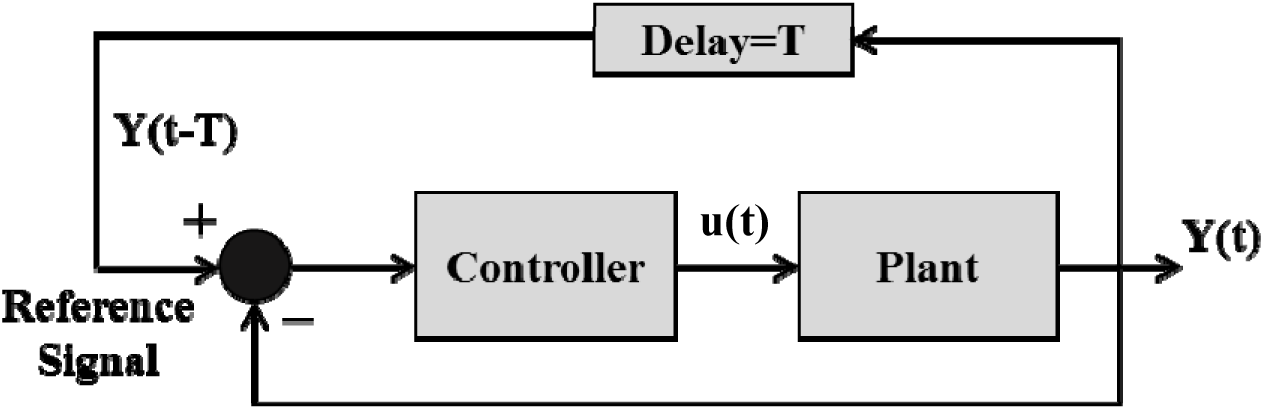
The components of the black box model with a variable reference signal designed based on time-delayed feedback control concepts.

According to Fig. 4 and setting the value of T on one, it can be said that the controller sends an optimum control signal to keep the output near its previous value. In this case, due to the limitation of the control signal, compensation and control purpose cannot be fully realized. However, in both conditions 2 and 3, the controller’s effort will partially prevent growth and increase of the output (i.e., RT) over time. In the results, it has been shown that which of these conditions is more suitable to avoid increasing the output (i.e., RT) over time.

### 2.2 Experiments Protocol

Three human experiments with different protocols were designed to compare and validate the computational outcomes under the mentioned three conditions. In these experiments, the performance of individuals was evaluated when 1-there was no feedback, 2-there was feedback with a fixed reference signal, and 3-feedback with a variable reference signal designed based on time-delayed feedback control concepts. The feedback was the difference between the value of RT in each trial and the reference signal. The value of this feedback was expressed in a fuzzy visual form to be understood by children. These feedbacks are described in the following paragraphs.

In all three conditions, participants performed a visual and auditory CPT. The main part of this CPT was composed of 400 trials. These 400 trials concluded the repetition of four blocks of 100 trials. In each block, numbers “1” and “2” (in Persian) were presented visually and auditory in a pseudo-random order. Participants were requested to press a bottom as soon as seeing or hearing the number “1” and do nothing for the number “2.” Each block had two parts: frequent and rare. In the frequent part, 84% of trials were the target (i.e., the number “1”). In the rare part, 84% of trials were the non-target (i.e., the number “2”). Visual stimuli (target or non-target) were presented for 170 milliseconds, and auditory ones were played for 500 milliseconds. The height of the visual stimuli was about 3.8 centimeters. The inter-stimulus interval was set at 1.8 seconds. Therefore, the main part of the task was 12 minutes. The stimuli were presented on a 21” LCD screen with the resolution of 1600×900 pixels and the refresh rate of 60 Hz. The distance of participants from the screen was about 70 cm. Auditory stimuli were played by a headphone (Somic® ST-1608) into the left and right ears. The accuracy and RT of responses were recorded during the task. All experiments were designed in Visual C# 2012 software running under Microsoft Windows 7 and controlled the presentation and storage procedures.

In the first experiment, participates received no feedback from their performance.

In the second experiment, the average RTs of 10 appropriate responses (i.e., response to target between 70 to 1100 milliseconds) from last trials of warming and practice phase was calculated before the start of the main phase. This average value was considered as the reference signal. In the main part, the individuals’ response time to the target was compared with the reference signal.

- If the individual’s response was correct (i.e., response to target) and his/her RT was equal or lower than the reference signal, then a full apple (score=1) was given to the subject as feedback.
- If the individual’s response was correct (i.e., response to target) but his/her RT was higher than the reference signal, then a half-eaten apple (score=0.5) was given to the subject as feedback.
- If the individual’s response was incorrect (i.e., response to non-target or no response to target), then a completely-eaten apple (i.e., score=0) was given to the subject as feedback.

Fig. 5 shows the image of the mentioned feedbacks. Participants were instructed about the task and the meaning of each feedback before the test. Fig. 6 demonstrates a view of one trial of the task with the presentation of feedback.

**Fig. 5.**
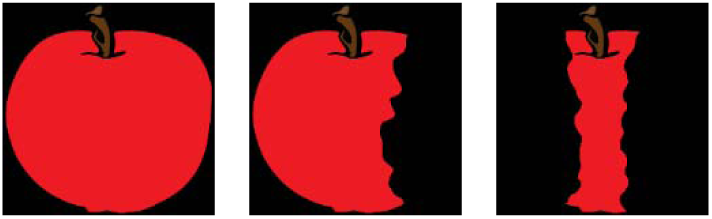
The image of feedbacks given to subjects during the task.

**Fig. 6.**
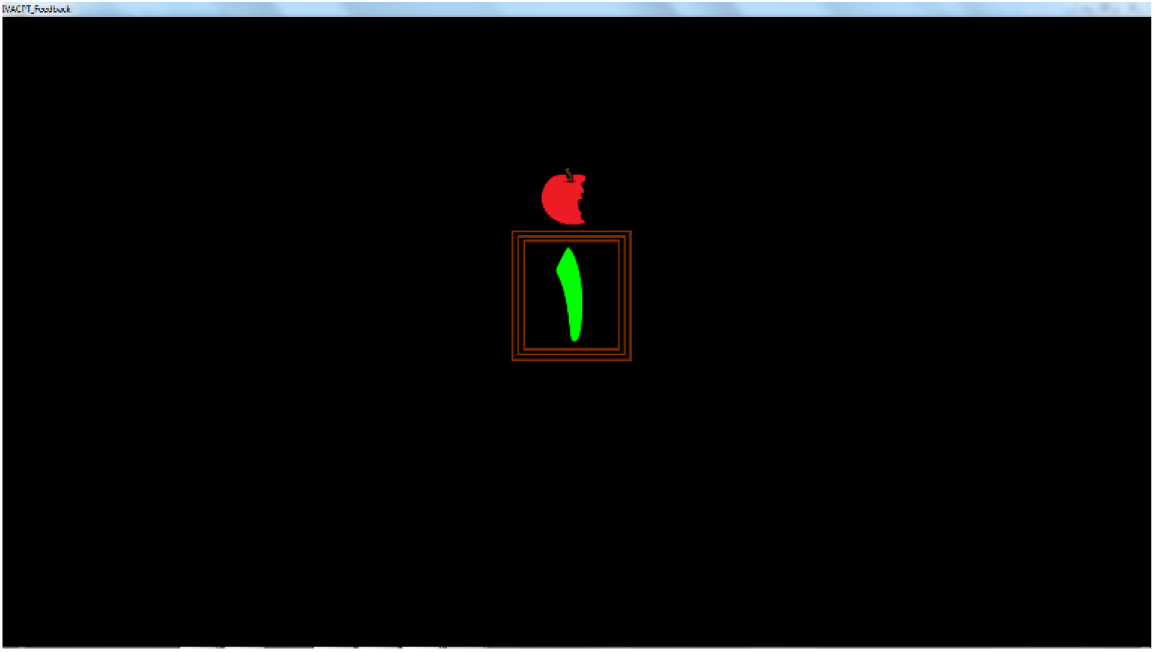
A view of one trial of the task with the presentation of the target (number “1” in Persian) and the feedback.

The third experiment was similar to the second one, with the difference that the reference signal was the correct RT of a previous trial (i.e., the last successful response to target). That is, the visual feedback was given based on the comparison between the current trial’s correct RT (i.e., response to target) and the correct RT in a previous trial (i.e., time-delayed feedback concept). The feedback was presented immediately after the participants’ response and remained for 200 msec on the screen. In trials with no response, the feedback was presented 1200 milliseconds after the presentation of the stimulus.

### 2.3 Participants

In the first experiment (without feedback), twelve typically developing children (9.2±0.6 years; one left-handed; eight girls) and eight children with ADD (9.6±0.8 years; two left-handed; three girls) participated.

In the second experiment (fixed reference feedback control), sixteen typically developing children (9.1±0.7 years; no left-handed; nine girls) and eight children with ADD (9.8±1 years; no left-handed; five girls) took part.

In the third experiment (time-delayed reference feedback control), twenty typically developing children (9.3±0.8 years; one left-handed; eleven girls) and eight children with ADD (9±0.5 years; one left-handed; three girls) participated. Eight children with ADD were not the same in three groups.

The experts in the ATIEH clinical neuroscience center based on DSM-IV criteria, an integrated visual and auditory CPT, and QEEG features made the diagnosis of children with ADD. Participants were not treated with any medication or behavioral interventions. For typically developing children, the National Institute for Children’s Health Quality (NICHQ) Vanderbilt assessment scales questionnaire was filled by the parents to ensure the absence of behavioral problems. The intelligence of all children was assessed based on the Goodenough–Harris test. All participants were classified into the “High average” or “High” class of IQ (e.g., a score of at least 110). They had no auditory problem or any otorhinolaryngological diseases, and they had normal vision.

All parents were aware of the protocol of the experiment and confirmed the informed consent form. All children participated voluntarily. Before the experiment, they were told that if they could complete the test, they would receive a prize. This sentence was only used to encourage them to engage more in the test, and at the end of the experiment, each of them received a gift. Iran University of Medical Sciences (#IR.IUMS.REC.1395 90133916) has approved the protocol of the study.

## 3. Results

Fig. 7 shows the output of the computational model (i.e., plant and controller) in three conditions: with no feedback, with fixed reference feedback, and with time-delayed reference feedback. The optimum control signal (*u*(*t*)) that was calculated during the optimization procedure to minimize the cost function, Eq. (2), was also demonstrated in Fig. 7 for both fixed and time-delayed references.

**Fig. 7.**
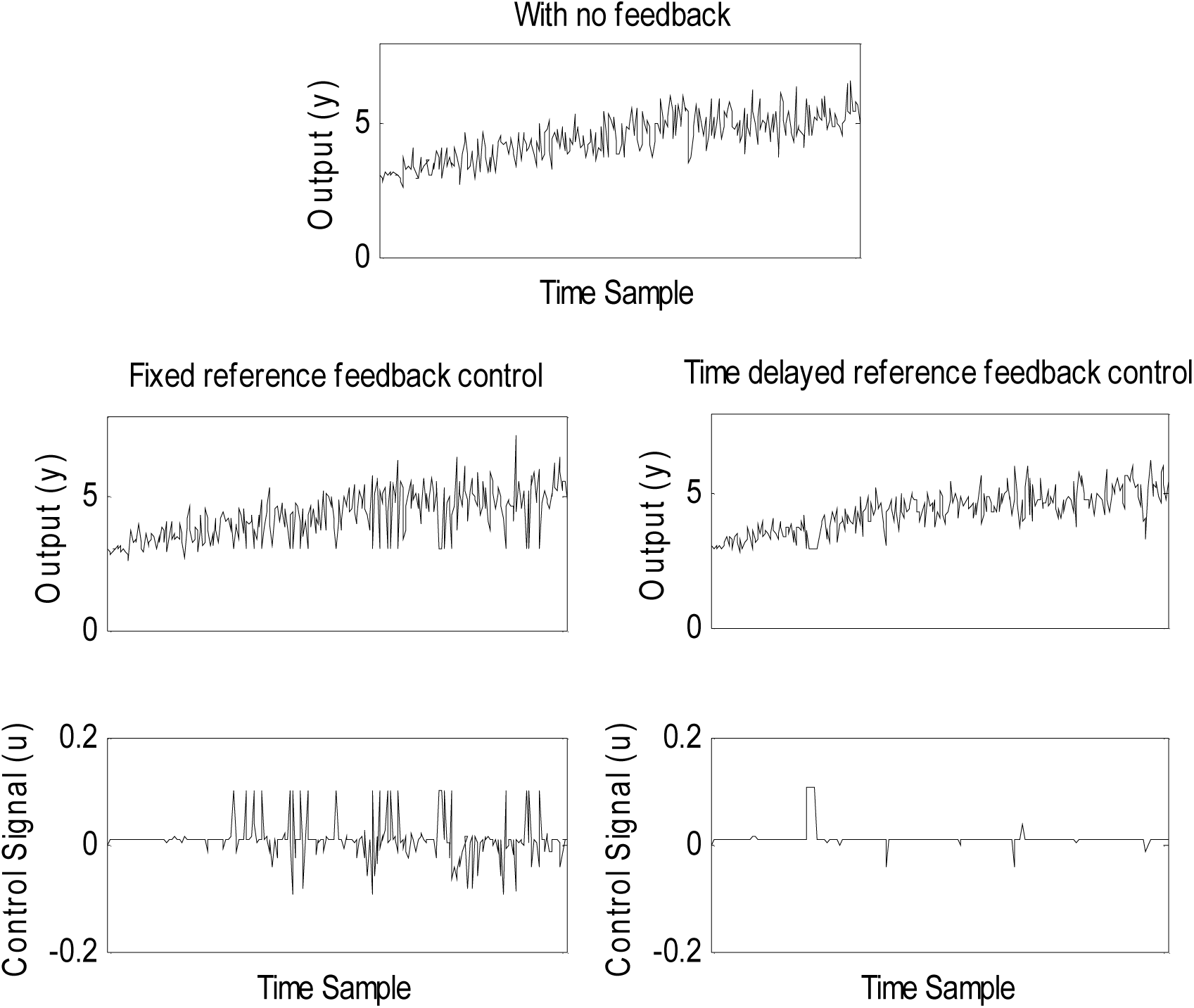
The output of the computational model (*Y*(*t*)) in different conditions: 1-with no feedback, 2-with fixed reference feedback, and 3-with time-delayed reference feedback. The last row shows the calculated optimum control signal (*u*(*t*)) in the second and third conditions

According to Fig. 7, the growth of output values in the time-delayed reference feedback is lower than the fixed reference. It can be observed that this stability obtained with lower values of the control signal. The norms of control signals in the fixed and time-delayed reference feedback are respectively 0.6 and 0.3.

Fig. 8 shows the correct RT signals recorded from typically developing and ADD groups in the mentioned three conditions.

**Fig. 8.**
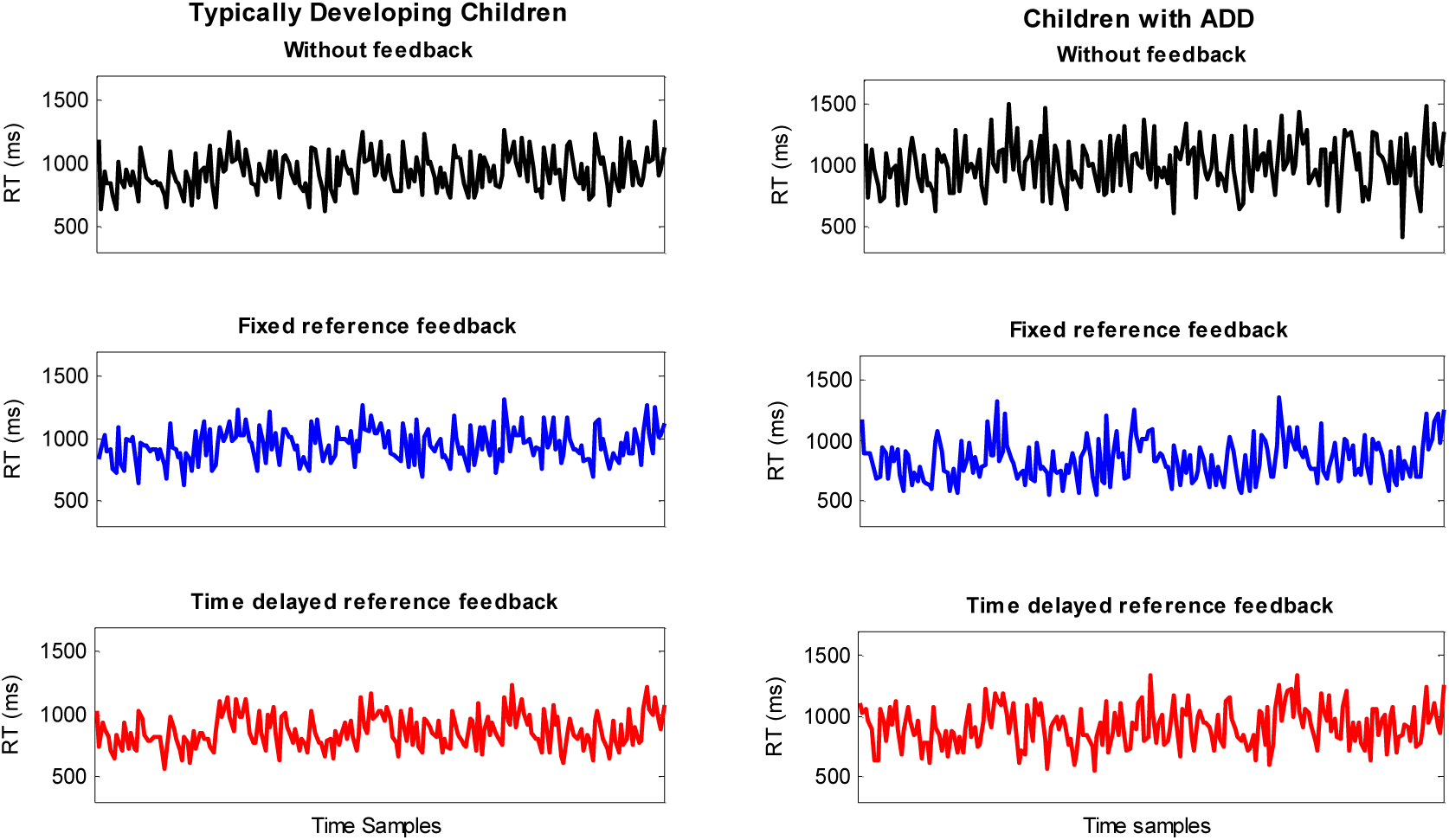
Signals of mean correct RTs recorded from typically developing and children with ADD in different conditions: 1-without feedback, 2-with fixed reference feedback, and 3-with a time-delayed reference feedback

According to Fig. 8, it can be observed that based on the recorded signals, one cannot be able to judge the differences between the three different situations, visually. Therefore, extracting an index from these signals seems necessary.

As mentioned before, the control object is to stabilize the RT value and to avoid its increment during a CPT task. Therefore, the ratio of mean correct RTs in the second half of the test into the first half was considered as an index to compare different conditions in the computational model and human experiments. Lower values of this ratio were interpreted as better performance. Figs. 9 and 10 show the value of this ratio in three conditions for the output of the computational model and human experiments.

**Fig. 9.**
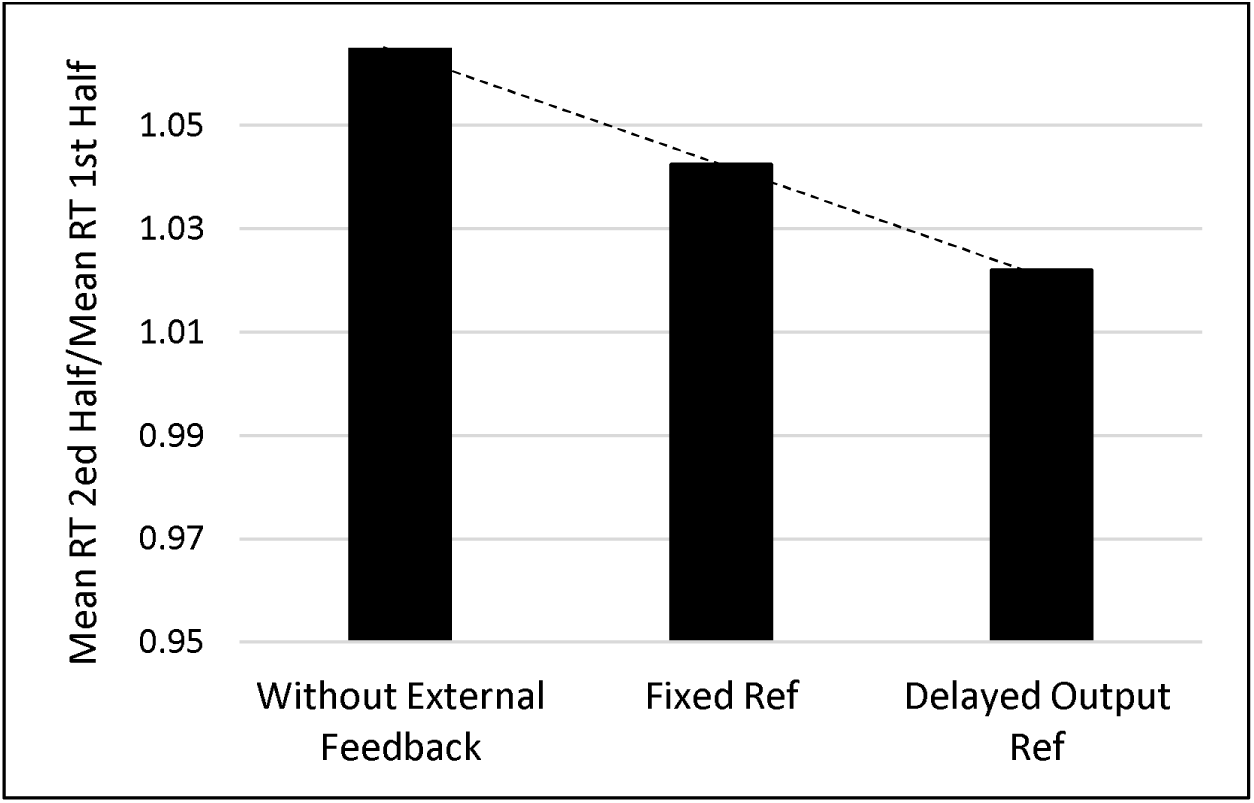
The effect of feedback with different reference signals on the ratio of the average of the model’s outputs (*Y*(*t*)) in several repetitions in the second half of the simulation time into the first.

**Fig. 10.**
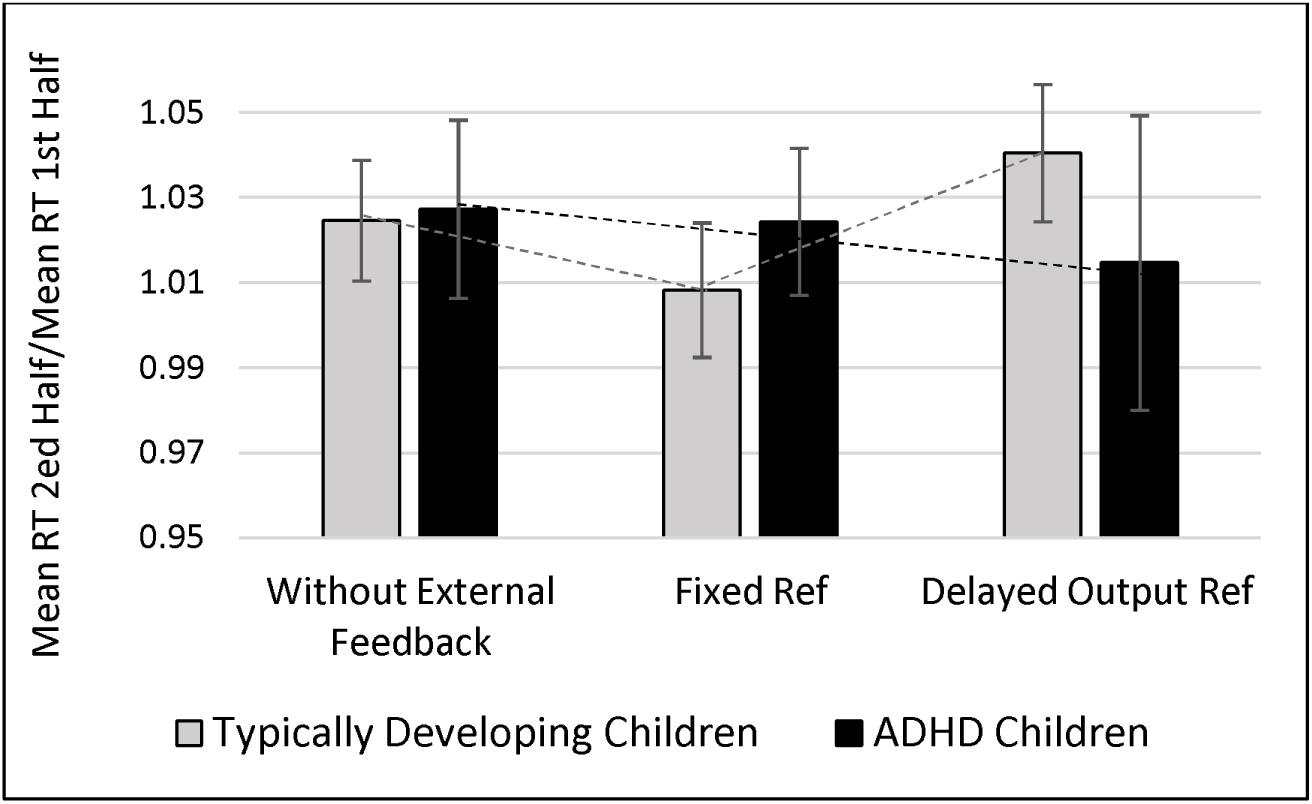
The effect of feedback with different reference signals on the ratio of the mean correct RTs in the second half of the test into the first in ADD and typically developing groups.

Two-way ANOVA with Sidak’s multiple comparison test showed that there is no statistical difference between three feedback conditions (F (2, 66) = 0.1789, p = 0.8366), between two groups of children (F (1, 66) = 0.0193, p = 0.89), and their interaction (F (2, 66) = 0.5685, p = 0.5691). However, some differences can be seen visually. According to Fig. 9, the output of the computational model shows that using the time-delayed feedback may lead to better performance than using the fixed reference feedback. This outcome is consistent with the results obtained from children with ADD (Fig. 10). However, in typically developing subjects, time-delayed feedback not only did not improve the performance relative to the fixed reference state but also led to worse performance than the state with no feedback.

## 4. Discussion

In neurocognitive rehabilitation methods, biofeedback is usually used as a complementary therapy. In this method, the performance of an individual is compared with an ideal standard reference. The result of this comparison is given to the individual as feedback. The individual’s neural controller uses this feedback to modify his/her performance [11]. The mentioned ideal standard performance, which is called the reference signal, is usually determined based on the performance of some ideal typically developing children or is measured from the individual when it is in an ideal condition. In both of these methods, the reference signal is fixed over time. In the current study, another method was proposed to define the reference signal. The time-delayed feedback control algorithm has inspired this method. In a computational view, feedback can deform the function of the system to a more stable condition. The controller can also compensate for the distance between the system’s output and the desired performance using the feedback error. Our computational and behavioral results are consistent with previous studies that showed the positive effects of biofeedback on human behavioral and physiological signals [25-27].

To examine the efficiency of the proposed reference signal, we compared the growth rate of RT in three conditions during a CPT. In the first condition, no feedback was given to the computational plant and the participants. In the second condition, the reference signal was the average of some correct RTs from the first trials of the test. It is believed that individual attention level in first trials is higher than the last. Therefore, this average value can be considered as a fix reference signal that relates to the ideal condition of the individual. The computational model showed that the fixed reference feedback control could improve the performance of the model concerning the state with no feedback. The outcomes obtained from the human experiment showed nearly the same result.

In the third condition, the reference signal was variable. It was equal to a correct RT in one step ago (i.e., time-delayed reference). The computational model demonstrated that this variable reference signal, which was designed based on time-delayed feedback control concepts, led to a better performance than the fixed reference. This result was congruent with the observation recorded from children with ADD. However, in typically developing children, a completely different result was observed. In typically developing children, time-delayed reference feedback control had a negative effect on the performance.

Regarding the explanation provided in previous sections, in time-delayed reference, the change of the response speed in comparison with a previous trial is given back to the subject. In a computational view, it seems like a first-order derivative in the control system. If the system is intrinsically stable, adding a derivative can cause the system becomes unstable. This result suggests that a difference between typically developing and children with ADD may exist in their performance monitoring system. The importance of self-monitoring capability was shown in the performance of individuals in cognitive tasks [42]. In previous studies, self-awareness and self-regulation problems have been reported in people with ADHD [43-45]. It can be claimed that typically developing children are aware of the changes in their performance continuously. Using this awareness, they can regulate their performance to some extents. It has been believed that the capability of internal monitoring can create an internal feedback loop that can affect performance and individual responses [46]. Therefore, when external information about the momentary alteration is provided for the system, it may affect the internal self-awareness. This effect may cause a conflict that can lead to worse performance. Thus, the impairment of brain regions involved in receiving, processing, and implementing momentary information about changes in the performance may lead to symptoms of inattention.

It is questionable that why did the performance drop not occurred in the fixed reference feedback condition in typically developing children. In the fixed reference feedback, the ongoing performance was compared with the first trials. Continues performance tests have a long duration. According to the storage limitation, remembering of the performance in the first trials is approximately impossible. Therefore, fixed reference feedback contains some new information for typically developing subjects that have no conflict with other internal information. However, this is a suggestion and needs more investigations.

As a result, in addition to the working memory capacity, feedback type, individuals’ ability, and task demand [29-32], the appropriate selection of the reference signal can affect the biofeedback result. According to the results of the current study, the reference signal should be selected based on the individual’s characteristics. In other words, it seems that biofeedback is sensitive to individual differences. This sensitivity reminds the single case methodologies, which have been recommended in neurobehavioural rehabilitation [47].

As shown in Fig. 7, the area under the control signal curve in the fixed reference feedback control is higher than that of in the time-delayed feedback control. The amount of mental effort in human experiments is represented by the control signal in the computational model. It that, the attention control system needs to consume energy to keep the performance (prevent an excessive rise in reaction time). Therefore, another prediction of the model is that performance improvement using time-delayed reference can be obtained with lower mental effort in comparison with a fixed reference signal in children with ADD.

## 5. Conclusion

As mentioned before, RT is an important index to quantify the performance of a subject. Therefore, to design a biofeedback system that works based on the RT changes, the reference signal can be fixed based on an ideal value or can be varied based on the concepts in the time-delayed control method. Results of this study showed that the selection of the reference signal is important in a biofeedback system. The incorrect selection of the reference signal can lead to more mental effort and energy consumption or worse performance. This study also suggests that the impairment of brain regions that process the information continuously may lead to the performance drop in a CPT. The development of new therapies to strengthen these centers can be effective in the cognitive rehabilitation of people with impaired attention, if this prediction is correct. However, increasing the number of participants and considering between-subjects variability can enhance the reliability of results. Examination of the effect of some other types of reference on the performance of the computational model and then the human subjects are also suggested for future works.

## Acknowledgments

This research has been supported by the cognitive sciences and technologies council grant (#201). We take this opportunity to thank the psychologists and psychiatrists of the ATIEH clinical neuroscience center for their kindly help in the procedure of the experiment and providing the required permission for carrying out the experiments. We would also like to thank the Amirkabir University of Technology for providing prizes that were given to the participants. Special thanks to the teachers of the Kowthar Primary for their help in arranging the experiments. We also thank the children’s parents for their patience during the experiment.

